# Aspergillus SUMOylation mutants have normal dynein function but exhibit chromatin bridges

**DOI:** 10.1101/2023.04.16.537086

**Authors:** Jun Zhang, Rongde Qiu, Baronger D. Bieger, C. Elizabeth Oakley, Berl R. Oakley, Martin J. Egan, Xin Xiang

## Abstract

Functions of protein SUMOylation remain incompletely understood in different cell types. The budding yeast SUMOylation machinery interacts with LIS1, a protein critical for dynein activation, but dynein-pathway components were not identified as SUMO-targets in the filamentous fungus *Aspergillus nidulans*. Via *A. nidulans* forward genetics, here we identified *ubaB*^Q247^*, a loss-of-function mutation in a SUMO-activation enzyme UbaB. Colonies of the *ubaB*^Q247^*, Δ*ubaB* and Δ*sumO* mutants looked similar and less healthy than the wild-type colony. In these mutants, about 10% of nuclei are connected by abnormal chromatin bridges, indicating the importance of SUMOylation in the completion of chromosome segregation. Nuclei connected by chromatin bridges are mostly in interphase, suggesting that these bridges do not prevent cell-cycle progression. UbaB-GFP localizes to interphase nuclei just like the previously studied SumO-GFP, but the nuclear signals disappear during mitosis when the nuclear pores are partially open, and the signals reappear after mitosis. The nuclear localization is consistent with many SUMO-targets being nuclear proteins, for example, topoisomerase II whose SUMOylation defect gives rise to chromatin bridges in mammalian cells. Unlike in mammalian cells, however, loss of SUMOylation in *A. nidulans* does not apparently affect the metaphase-to-anaphase transition, further highlighting differences in the requirements of SUMOylation in different cell types. Finally, loss of UbaB or SumO does not affect dynein-and LIS1-mediated early-endosome transport, indicating that SUMOylation is unnecessary for dynein or LIS1 function in *A. nidulans*.

## Introduction

Post-translational modification of proteins by SUMO (Small ubiquitin-like modifier) has been linked to many cellular processes, but the functions of SUMOylation in different cell types still need to be better understood (Vertegaal, 2022). SUMOylation has recently been implicated in regulating cytoskeletal proteins or proteins involved in intracellular motility (Alonso et al., 2015; Greenlee et al., 2018; Leisner et al., 2008; Meednu et al., 2008). Importantly, the budding yeast (*Saccharomyces cerevisiae*) homolog of LIS1, which is a key dynein activator (Markus et al., 2020), has been shown to physically interact with the SUMOylation machinery (Alonso et al., 2012). However, LIS1 or proteins in the dynein pathway were not among the previously identified Sumo targets in the filamentous fungus *Aspergillus nidulans* (Harting et al., 2013; Horio et al., 2019). Despite this difference, it would be useful to test directly whether SUMOylation affects dynein function in different organisms and cell types.

SUMOylation has been known to affect mitosis in various ways (Dasso, 2008; Dawlaty et al., 2008; Eifler et al., 2018; Eifler and Vertegaal, 2015; Johansson et al., 2021; Liu et al., 2023; Ptak et al., 2021; Sridharan and Azuma, 2016; Yatskevich et al., 2021). During a normal mitosis, replicated sister chromatids segregate and move to the opposite poles of the mitotic spindle during anaphase, which is followed by spindle disassembly and cytokinesis that divides the daughter cells. Chromatin bridges, a type of anaphase bridges, are caused by a failure in timely resolving of the the linkage between the duplicated sister chromatids (Finardi et al., 2020). Such a failure can be caused by many factors, including, but not limited to, dicentric chromosomes caused by abnormal fusions between telomeres or broken chromosome ends, and defects in resolving DNA catenanes caused by a topoisomerase II dysfunction or other problems (Dawlaty et al., 2008; Finardi et al., 2020; Flynn et al., 2021; Maciejowski et al., 2015; Nielsen et al., 2015; Umbreit et al., 2020). Most relevant to our current study, a defect in topoisomerase II SUMOylation has been shown to cause chromatin bridges (Dawlaty et al., 2008). Chromatin bridges have been linked to chromothripsis (Maciejowski et al., 2015; Umbreit et al., 2020), which causes dramatic rearrangement of one or a few chromosomes (Leibowitz et al., 2015; Ly and Cleveland, 2017). Chromatin bridges induced by some anti-mitotic drugs may also activate the cGAS-mediated type 1 interferon signaling (Flynn et al., 2021), which is a response to cytoplasmic DNA (Li and Chen, 2018). Whether the bridges affect cytokinesis, especially abscission, has been studied in *S. cerevisiae* and mammalian cells (Amaral et al., 2016; Carlton et al., 2012; Finardi et al., 2020; Nähse et al., 2017; Petsalaki and Zachos, 2021; Shi and King, 2005; Steigemann et al., 2009), where a NoCut pathway or abscission checkpoint is implicated in abscission delay in response to chromatin bridges (Amaral et al., 2016; Carlton et al., 2012; Norden et al., 2006; Steigemann et al., 2009). However, this does not prevent the breakage of some chromatin bridges via different mechanisms, which significantly enhances the genome instability observed in cancer cells (Maciejowski et al., 2015; Umbreit et al., 2020).

Many filamentous fungi contain hyphal syncytia with multiple well-spaced nuclei that are distributed by cytoplasmic dynein and other mechanisms (Mela et al., 2020; Xiang, 2018), and the presence of multiple nuclei in the same compartment should allow errors of chromosome segregation to be tolerated since the nuclei can complement each other for functions. The filamentous fungus *Aspergillus nidulans* is an established model organism for studying the regulation of cell cycle progression, among other interesting topics (De Souza et al., 2009; Dörter and Momany, 2016; Edgerton et al., 2015; Edgerton-Morgan and Oakley, 2012; Etxebeste and Espeso, 2020; Harris, 2001; Morris, 1975; Morris and Enos, 1992; Nayak et al., 2010; Osmani et al., 2006; Osmani and Mirabito, 2004; Paolillo et al., 2018; Suresh and Osmani, 2019). *A. nidulans* cells undergo a “semi-open” mitosis with partial disassembly of the nuclear pore complex during mitosis (De Souza et al., 2004), a mode that differs from the more closed and more open modes of mitosis in yeast and mammalian cells (De Souza and Osmani, 2009; Dey and Baum, 2021). Unlike another filamentous fungus *Ashbya gossypii* where each nucleus undergoes mitosis autonomously within a syncytium (Gladfelter et al., 2006; Roberts and Gladfelter, 2015), nuclei within a syncytium of *A. nidulans* undergo mitosis almost synchronously (Rosenberger and Kessel, 1967; Suelmann et al., 1997). Syncytia are separated by a cross wall-like structure called the septum, whose formation is triggered by the septation-initiation network (SIN), but it does not necessarily couple with nuclear division, which differs from septation/cytokinesis in yeasts (Kim et al., 2006; Kim et al., 2009; Krapp et al., 2004; Wolfe and Gould, 2005). In filamentous fungi, each septum contains a septal pore, which allows some intercompartmental exchanges during interphase, and the septal pore in *A. nidulans* closes during mitosis using a NimA-kinase-involved mechanism (Shen et al., 2014), which differs from the Woronin-body-mediated septal-pore-sealing in response to hyphal injury (Jedd and Chua, 2000; Mamun et al., 2023; Shen et al., 2014; Songster et al., 2023; Steinberg et al., 2017; Tenney et al., 2000). It is not clear whether chromatin bridges prevent the sealing of septal pores during mitosis in *A. nidulans*.

In this current study, we obtained an *A. nidulans* mutant exhibiting chromatin bridges and identified the causal mutation in a SUMO-activating enzyme UbaB. Our data suggest that SUMOylation in *A. nidulans* is important for resolving chromatin bridges during mitosis, but it is not critical for LIS1 or dynein function in *A. nidulans*. Our observations on the chromatin bridges also suggest that the bridges prevent neither cell cycle progression nor the mitotic septal pore sealing in *A. nidulans*.

## Results and Discussion

### A loss-of-function mutation in a sumo-activating enzyme UbaB produces abnormal chromatin bridges

From a UV mutagenesis, we obtained a mutant exhibiting an apparently increased number of bi-nuclei pairs, some of which are connected by chromatin bridges (Figure 1A), as judged by GFP-labeled Histone H1 (Xiong and Oakley, 2009). Bi-nuclei pairs can also be observed in wild-type cells as they represent chromosomes separated after anaphase. However, chromatin bridges are rarely observed in a wild-type control strain, and persistent chromatin bridges are consistent with a defect in segregating the duplicated chromosomes during anaphase (Finardi et al., 2020). To identify the causal mutation, we took an approach combining *A. nidulans* genetics, whole genome sequencing and bioinformatic analysis (see Materials and Methods for more details). Using this approach, we identified the causal mutation in AN2450 (using the fungiDB.org gene designation), which encodes UbaB, the key component of the sole SUMO-activating enzyme in *A. nidulans* (Harting et al., 2013). UbaB has 610 amino acids, and we identified a C to T mutation, that changed the codon CAA (Glutamine or Q) at aa247 to the stop codon TAA, thereby removing a big portion of UbaB. We named the mutant *ubaB*^Q247^*.

**Figure 1.**
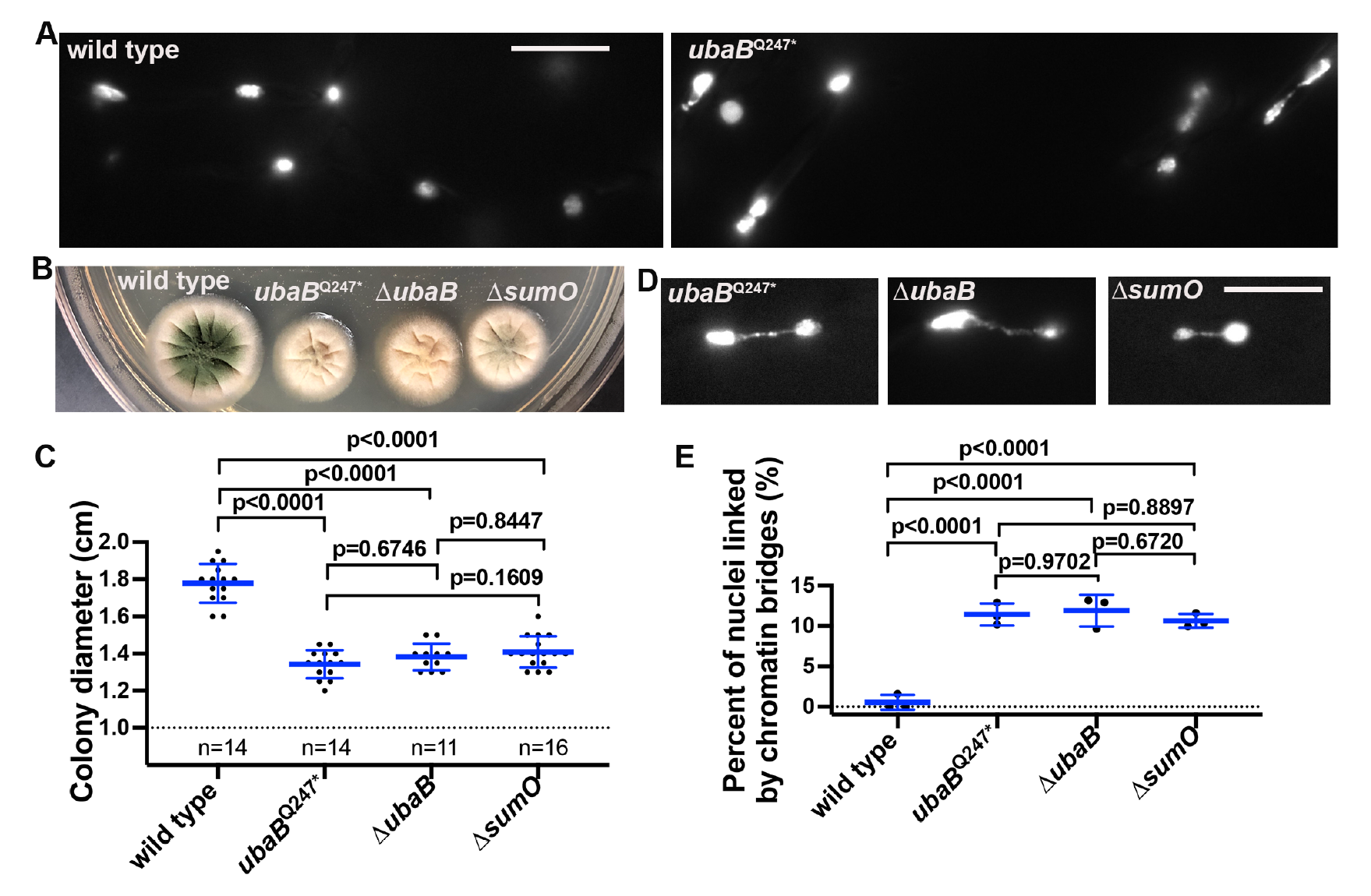
Colony and chromatin-bridge phenotypes of the *ubaB*^Q247^*, Δ*ubaB* and Δ*sumO* mutants. (A) The nuclear phenotype of the *ubaB*^Q247^* mutant visualized by using a Histone-H1- GFP fusion. Bar, 10 μm. (B) Colony phenotypes of wild-type, the *ubaB*^Q247^*, Δ*ubaB* and Δ*sumO* strains. (C) A quantitative analysis of colony diameter of wild-type, the *ubaB*^Q247^*, Δ*ubaB* and Δ*sumO* strains (Ordinary one-way ANOVA with Tukey’s multiple comparisons test). Scatter plots with mean and S.D. values were generated by Prism 9. (D) Chromatin bridges in the *ubaB*^Q247^*, Δ*ubaB* and Δ*sumO* strains containing Histone-H1-GFP. Bar, 10 μm. (E) A quantitative analysis on the percent of nuclei linked by chromatin bridges in the wild-type, the *ubaB*^Q247^*, Δ*ubaB* and Δ*sumO* strains (Ordinary one-way ANOVA with Tukey’s multiple comparisons test). Scatter plots with mean and S.D. values were generated by Prism 9.

The *S. cerevisiae* ortholog of UbaB, Uba2, is essential for colony growth (Dohmen et al., 1995; Johnson et al., 1997). However, the *A. nidulans ubaB*^Q247^* mutant forms a colony, albeit smaller than that of a wild type (Figure 1B, 1C). In *S. cerevisiae* and other organisms, Aos1 forms a heterodimer with Uba2/UbaB, which plays a critical role in SUMO-activation (Johnson et al., 1997; Shih et al., 2002). In *S. cerevisiae*, Aos1 (activation of Smt3p) is also essential (Johnson et al., 1997). In addition, the only SUMO-encoding gene Smt3p in *S. cerevisiae* is also essential, while Pmt3p, its ortholog in the fission yeast *Schizzosaccharomyces pombe*, is not, but it plays a critical role in growth as it affects multiple processes including telomere length control and chromosome segregation (Tanaka et al., 1999).

To further address the cellular function of UbaB in *A. nidulans*, we constructed a Δ*ubaB* mutant in which the coding region of the *ubaB* gene is deleted (Figure S1). We analyzed the phenotypes of this Δ*ubaB* mutant together with a previously constructed Δ*sumO* mutant in which the only SUMO-encoding gene in *A. nidulans* is deleted (Wong et al., 2008). The *ubaB*^Q247^*, Δ*ubaB* and Δ*sumO* mutants produced similar colonies that are smaller than the wild-type colony (Figure 1B, 1C), and they also appeared to have reduced conidiation as shown previously (note that another Δ*ubaB* mutant was constructed previously) (Horio et al., 2019; Wong et al., 2008). Importantly, the Δ*ubaB* and Δ*sumO* mutants also exhibit chromatin bridges similar to those exhibited in the *ubaB*^Q247^* mutant (Figure 1D). In all three of these mutants, ∼10% of the total nuclei are connected by visible chromatin bridges, a percentage that is significantly higher than that in wild type but not significantly different among the three mutants themselves (Figure 1E). These data indicate that *ubaB*^Q247^* is a loss-of-function mutation, and that UbaB-mediated SUMOylation is important for the completion of chromosome segregation in *A. nidulans*.

### Chromatin bridges do not prevent cell cycle progression or mitotic septal-pore sealing

To examine the effects of chromatin bridges on cell cycle progression, we observed both the *ubaB*^Q247^* mutant and the Δ*ubaB* mutant containing Histone-H1-GFP (Xiong and Oakley, 2009) and the NLS-DsRed fusion that labels the interphase nuclei (De Souza et al., 2004; Suelmann et al., 1997). Previous work has shown that the NLS-DsRed fusion leaks out of the nuclei to become cytoplasmic due to the partial disassembly of the nuclear pore complex (De Souza et al., 2004). Importantly, the position of a sealed septum at mitosis can be inferred by the boundary of the NLS-DsRed signals (Shen et al., 2014). In our wild-type control strain, we were able to see the diffused NLS-DsRed signals during mitosis and the mitotic septal-pore sealing just as described previously (Figure S2) (Shen et al., 2014). In the Δ*ubaB* mutant, the vast majority of chromatin bridges appeared to be at interphase; specifically, 97% of randomly chosen Histone-H1-GFP-labeled bridges (n=33) were surrounded by nuclear NLS-DsRed signals (Figure 2A, Figure S3). Furthermore, we observed Δ*ubaB* cells containing GFP-tubA that labels microtubules (GFP-microtubules) (Horio and Oakley, 2005; Xiong and Oakley, 2009). We found that most hyphae containing the NLS-DsRed-labeled bridges (∼85%, n=27) exhibit cytoplasmic microtubules typical of interphase cells, rather than mitotic spindles (Figure 2B, Figure S4), indicating that bridge persistence is not causally linked to a hyperstability of spindle microtubules in these mutants. One time-lapse series of the *ubaB*^Q247^* mutant also shows that interphase nuclei connected by a chromatin bridge can re-enter mitosis and then exit mitosis, as judged by the disappearance and reappearance of NLS-DsRed signals (Figure 3, Video 1), although the exit from mitosis in this case happened abnormally without nuclear division. Together, all these results suggest that the chromatin bridges do not prevent cell cycle progression. In addition, the presence of the NLS-DsRed signals in the bridges and connected nuclei suggest that the bridges do not grossly destroy the integrity of nuclear membrane and nuclear pore complexes.

**Figure 2.**
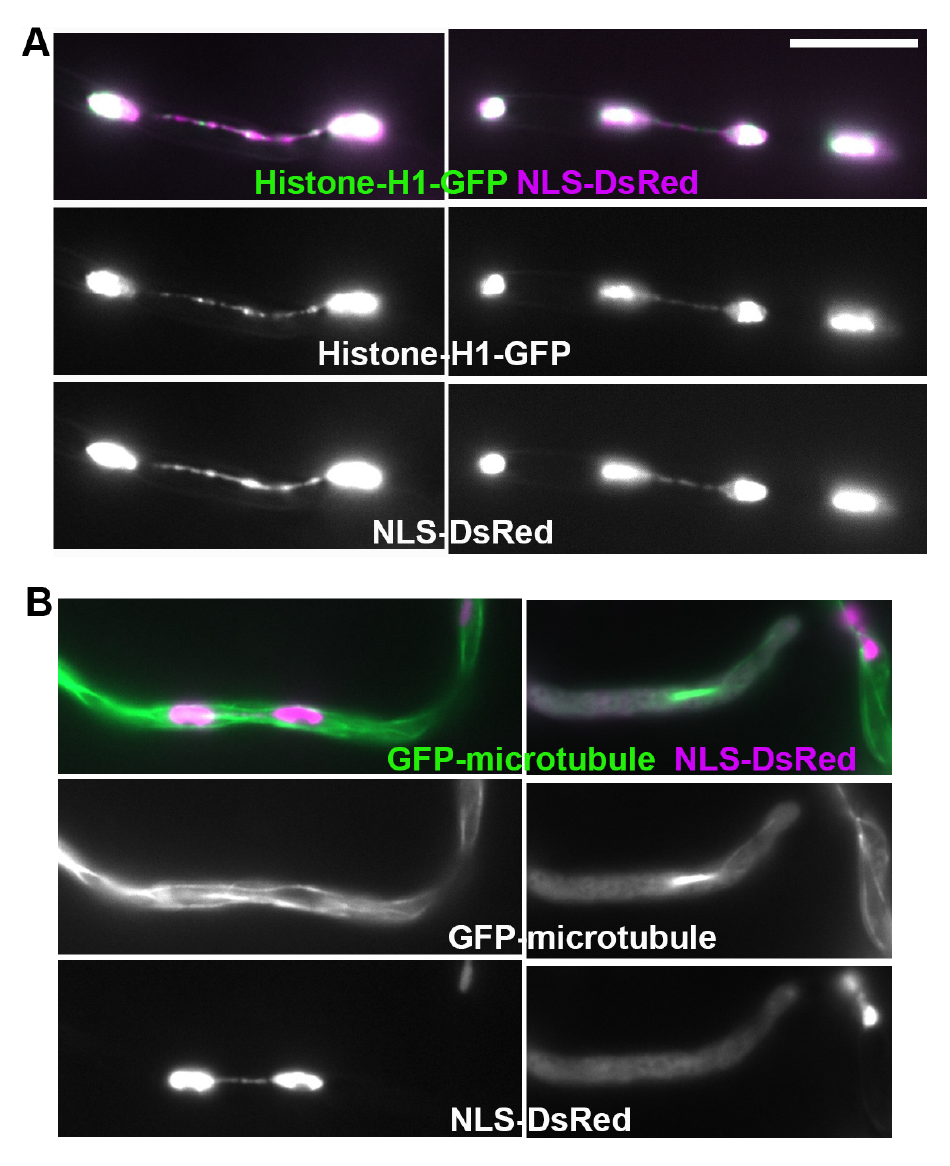
Images of the Δ*ubaB* mutant showing that chromatin bridges can persist through the mitosis/interphase transition and are not linked to hyperstability of mitotic spindles. (A) Images of the Δ*ubaB* mutant showing that nuclei connected by chromatin bridges can enter interphase as evidenced by the nuclear NLS-DsRed signals. Bar, 10 μm. (B) Images on the left show a NLS-DsRed-labeled bridge surrounded by cytoplasmic microtubules typical of interphase cells. A mitotic cell containing spindle microtubules is shown on the right for comparison. Note that the NLS-DsRed signals are diffused in the cytoplasm during mitosis.

**Figure 3.**
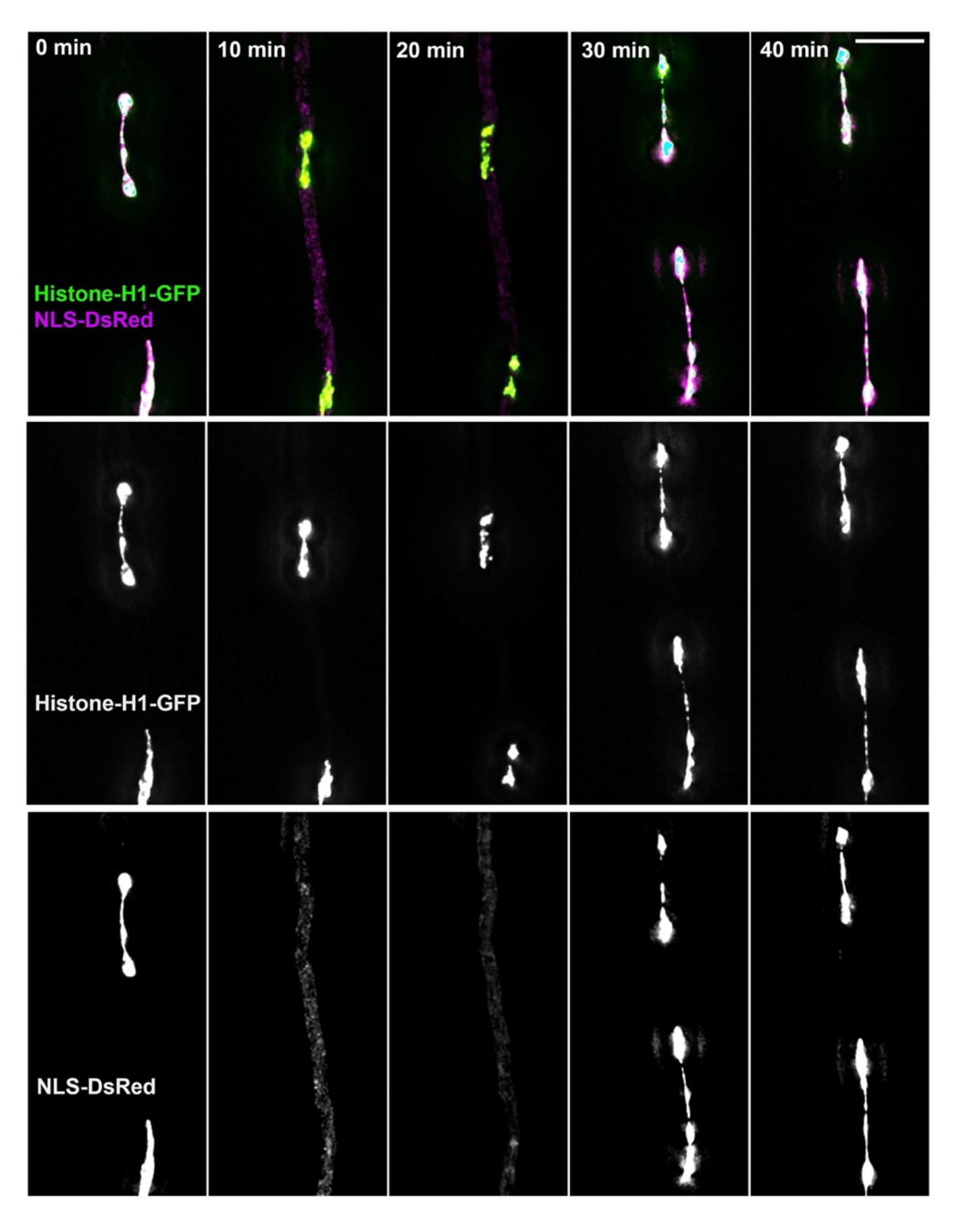
Interphase nuclei connected by chromatin bridges can enter mitosis and exit from mitosis without resolving the bridges. These images of the *ubaB*^Q247^* mutant are from a time-lapse sequence (taken at 10-min intervals) shown in Video 1. Note that entry into mitosis (10 min) is indicated by the disappearance of NLS-DsRed signals from the nuclei and exit from mitosis (30 min) is indicated by the reappearance of NLS-DsRed signals in the nuclei. Bar, 10 μm.

Our finding that chromatin bridges do not prevent cell cycle progression is consistent with observations made in mammalian cells (Maciejowski et al., 2015; Steigemann et al., 2009; Umbreit et al., 2020), although our results do not suggest any obvious nuclear membrane damage as observed in mammalian cells (Maciejowski et al., 2015). In both *S. cerevisiae* and mammalian cells, an Aurora-B-kinase-mediated NoCut pathway or abscission checkpoint has been proposed to prevent premature cutting of DNAs in the chromatin bridges (Carlton et al., 2012; Norden et al., 2006; Steigemann et al., 2009) (although Aurora B inhibition in the presence of bridges prevents furrow regression rather than abscission (Steigemann et al., 2009)). Interestingly, we have detected a septum on a chromatin bridge during mitosis, as judged by the appearance of cytoplasmic NLS-DsRed signals only at one side of the septum (Figure 4; Video 1) (Shen et al., 2014). That the NLS-DsRed signals are only present at one side is a strong indication that the septum pore is sealed (Shen et al., 2014). This is a rare event caught by serendipity (because mitotic cells are very rare (<5%) in a cell population and most chromatin bridges are present during interphase), but it clearly argues against the presence of an effective checkpoint that would prevent bridge cutting in these mutants. We would not rule out the possibility that loss of SUMOylation affects a possible checkpoint, especially since Aurora-B can be sumoylated (Ban et al., 2011; Fernández-Miranda et al., 2010). However, Aurora-B SUMOylation has never been implicated in the abscission checkpoint, and the only *A. nidulans* Aurora kinase (De Souza et al., 2017) has never been identified as a sumo target (Harting et al., 2013; Horio et al., 2019). In *A. nidulans*, although cytokinesis (or septation) is not coupled to nuclear division, and septal-pore sealing happens during mitosis rather than after nuclear division (Shen et al., 2014), Aurora kinase does localize to chromatin bridges that were caused by a deletion of core Nups of the nuclear pore complex (Chemudupati et al., 2019). Importantly, these bridges are linked to a hyperstability of mitotic spindles (Chemudupati et al., 2019). In *S. cerevisiae*, hyperstability of mitotic spindles has been causally linked to Aurora-B-mediated bridge monitoring (Amaral et al., 2016), suggesting that it is an alteration of the spindle rather than a chromatin bridge *per se* that is being monitored. Since the chromatin bridges in the *ubaB* mutants are mainly inside interphase cells, they may not be monitored by any effective checkpoint.

**Figure 4.**
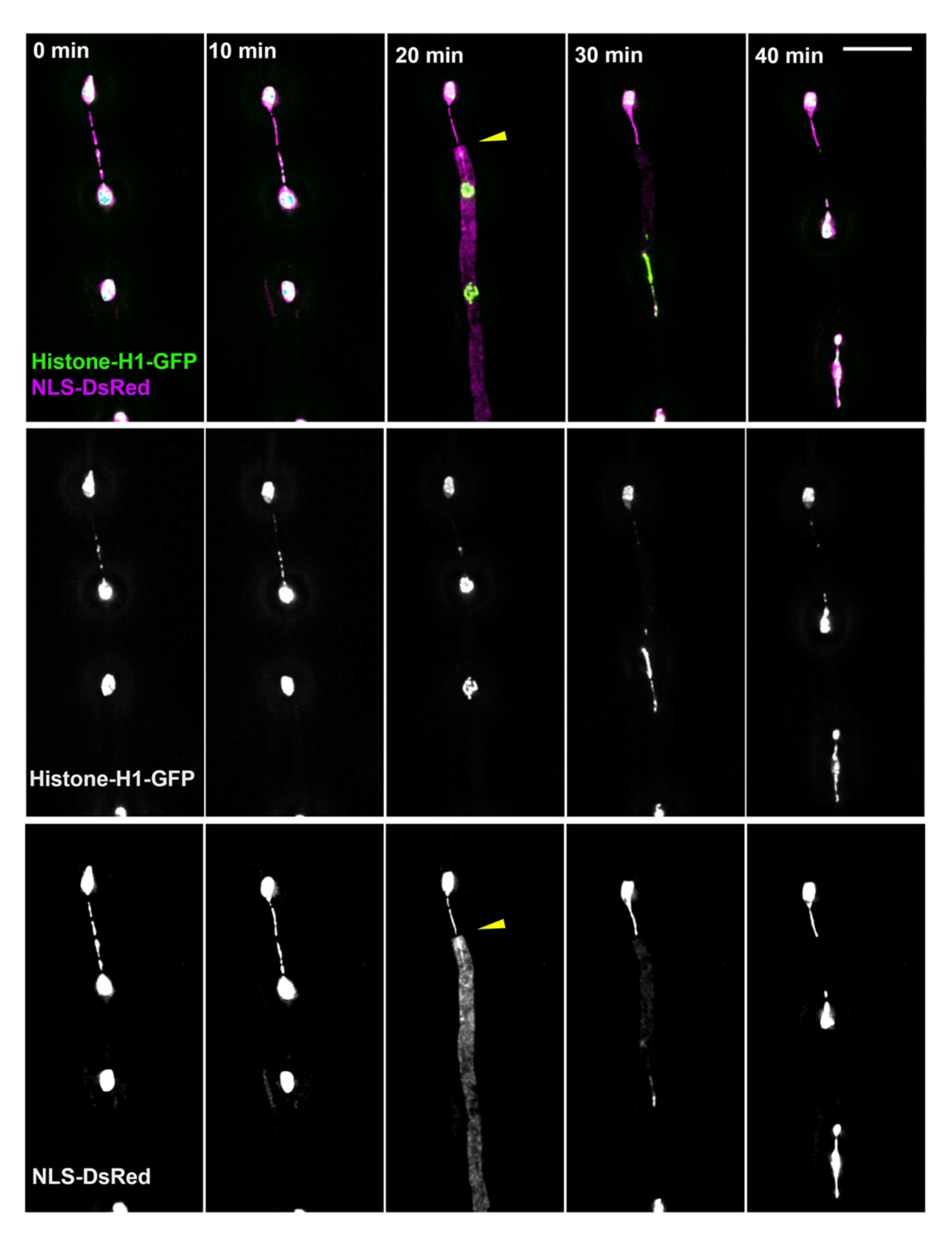
Septum formation and septal pore closure on a chromatin bridge. These images of the *ubaB*^Q247^* mutant are from a time-lapse sequence (taken at 10-min intervals) shown in Video 1. The two top nuclei were connected by a chromatin bridge at time points 0 min and 10 min. The presence of a closed septum (indicated by a yellow arrowhead) was evidenced by the appearance of NLS-DsRed signals only on the lower compartment at the 20-min time point. Note that the septum might have formed before this time point during interphase, but the cytoplasmic NLS-DsRed signals during mitosis and the previous result that septal pores get closed during mitosis helped reveal the presence of the septum (Shen et al., 2014). Bar, 10 μm.

### The SUMO-activating enzyme UbaB localizes to nuclei during interphase

To observe UbaB localization, we constructed a strain containing a *ubaB*-GFP fusion gene integrated at the *ubaB* locus. UbaB-GFP localized to the nucleus in interphase cells (Figure 5). However, UbaB-GFP signals disappeared during mitosis, and only reappeared after the formation of daughter nuclei when the NLS-DsRed signals also reappeared (Figure 5). This localization pattern is similar to the previously studied SumO-GFP (Wong et al., 2008) and is consistent with many SUMO-targets in *A. nidulans* being nuclear proteins, including topoisomerase II (Horio et al., 2019). Because a defect in SUMOylation of topoisomerase II has been implicated in chromatin bridges in mammalian cells (Dawlaty et al., 2008), the chromatin-bridge phenotype of our *ubaB* mutants could be explained by compromised topoisomerase II activity that leads to defect in resolving DNA catenanes. That UbaB-GFP appears quickly in the daughter nuclei upon post-mitotic assembly of the nuclear pore complexes (De Souza et al., 2004) suggests that the proteins are imported robustly into the nuclei from the cytoplasm after mitosis.

**Figure 5.**
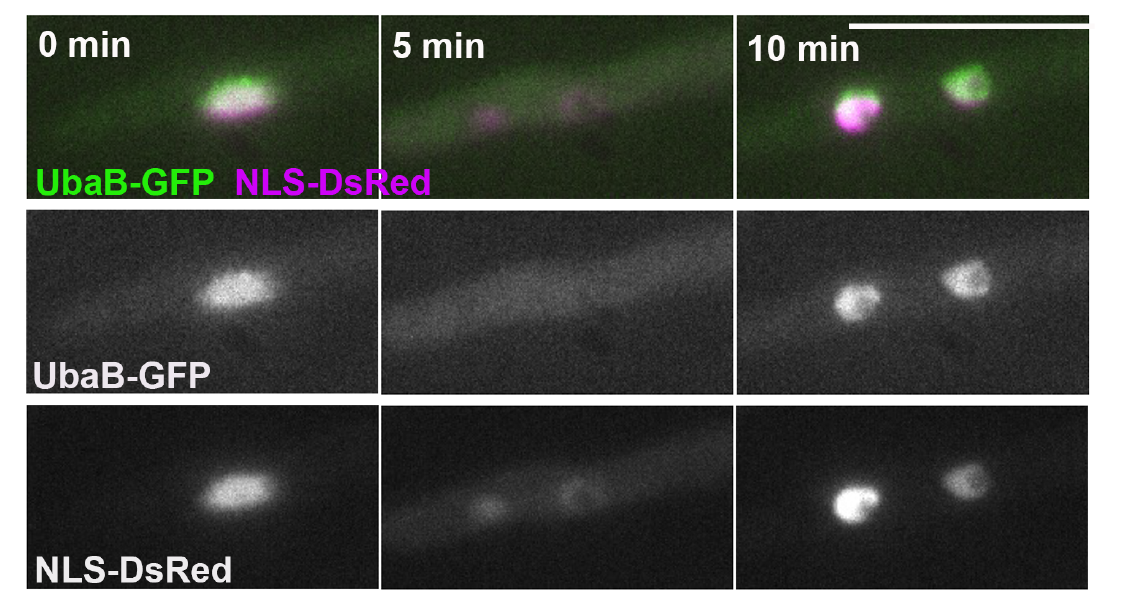
The nuclear localization of UbaB-GFP and its disappearance during mitosis. At 0 min, the germling contains one nucleus, and the UbaB-GFP is localized to the nucleus that is also labeled by NLS-DsRed. At the 5-min time point, UbaB-GFP nuclear signals disappeared. This is related to mitosis as evidenced by the faint nuclear signals of NLS-DsRed in two daughter nuclei. UbaB-GFP signals reappeared inside the two nuclei at 10 min, after the nuclear pore complex fully assembled as evidenced by the NLS-Ds-Rad signals. Bar, 10 μm.

### Loss of SUMOylation does not apparently affect the metaphase-to-anaphase transition in *A. nidulans*

It has been reported that SUMOylation of the anaphase-promoting complex/cyclosome (APC/C) plays an important role in the metaphase-to-anaphase transition in human cells, although the yeast APC/C subunits were not identified as sumo-targets (Eifler et al., 2018). In *A. nidulans*, none of the APC/C subunits have been identified as SUMO-targets (Harting et al., 2013; Horio et al., 2019). To address any possible role of SUMOylation in the metaphase-to-anaphase transition, we used the temperature-sensitive *nimT*23 (cdc25) mutant that is blocked in late G2 at the restrictive temperature of 42°C (O’Connell et al., 1992) and compared the *nimT*23 single mutant with a Δ*ubaB*, *nimT*23 double mutant after shifting cells to the permissive temperature of 25°C. As expected, we found that germ tubes produced after a 6-hour incubation at 42°C all contained a single G2 nucleus as shown by the Histone-H1-GFP. While the single nucleus in some germ tubes entered the germ tube as shown previously (Osmani et al., 1990; Xiang et al., 1994), about 80% of the *nimT*23 germ tubes (n=54) contained a single nucleus inside the spore head (Figure 6). Possibly, an interphase nucleus may move but not as robustly as the mitotic spindle at this early stage of fungal growth. This notion will be examined more carefully in the future, particularly in light of dynein-mediated movements of interphase nuclei and mitotic spindles in budding yeast (note that a spindle is contained inside the nucleus in yeast) (Winey and Bloom, 2012; Yeh et al., 1995).

**Figure 6.**
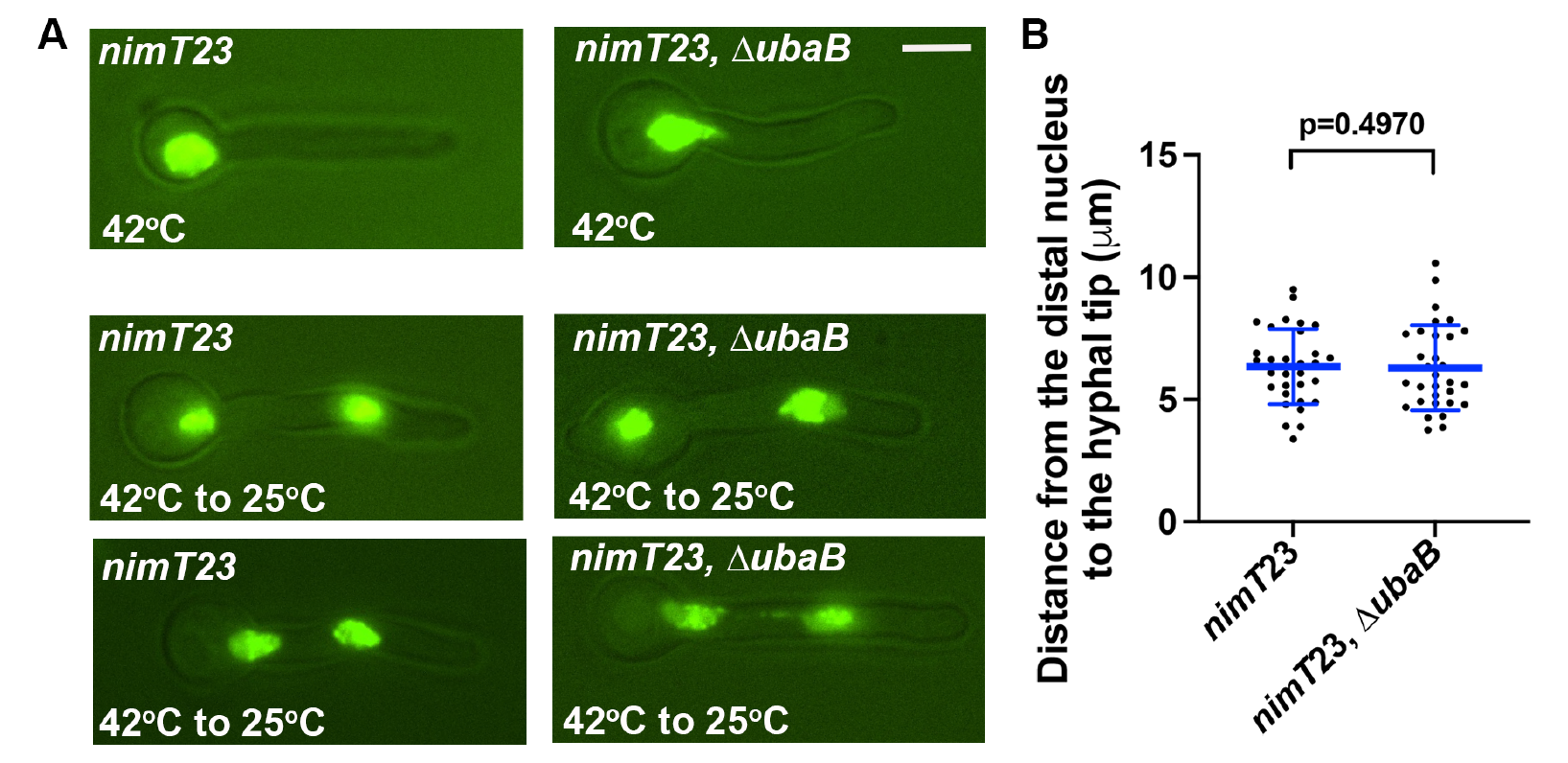
Loss of SUMOylation does not affect nuclear division and distribution in the *nimT*23 mutant that was initially blocked at G2. (A) Nuclei labeled with Histone H1-GFP in the *nimT*23 single mutant and the *nimT*23, Δ*ubaB* double mutant. After a 6-hr incubation at the restrictive temperature of 42°C, both mutants show a single nucleus. After shifting the cells to 25°C for 0.5- hr to 1-hr (42°C to 25°C), more than 90% of the germ tubes in both mutants show two nuclei. Bar, 5 μm. (B) An analysis on the distance between the distal nucleus and the hyphal tip revealed no significant difference between the two mutants (unpaired *t* test, two-tailed, n=31 for both mutants, and the data for both mutants passed the D’Agostino & Pearson normality test). Scatter plots with mean and S.D. values were generated by Prism 9.

Remarkably, images taken within 0.5-hr to 1-hr after shifting the *nimT*23 cells to room temperature (25°C) showed two nuclei in most germ tubes regardless of the Δ*ubaB* allele (96.7% in the *nimT*23 mutant, n=398; 94.9% in the Δ*ubaB*, *nimT*23 double mutant, n=234, note that only germ tubes that reached a length of ∼8 μm were counted). Thus, loss of SUMOylation does not produce an obvious defect in the metaphase-to-anaphase transition. This result further highlights the notion that sumo-targets and/or the effects of SUMOylation may vary in different organisms and cell types.

### Loss of SUMOylation does not apparently affect dynein function in *A. nidulans*

As mentioned earlier, within 0.5-hr to 1-hr after shifting the *nimT*23 single mutant and the Δ*ubaB*, *nimT*23 double mutant to room temperature, two nuclei were present in most germlings. Importantly, these nuclei were positioned properly regardless of the Δ*ubaB* allele (Figure 6), suggesting that loss of SUMOylation does not apparently affect dynein-mediated nuclear migration. In filamentous fungi, dynein and its activator LIS1 are critical not only for nuclear distribution but also for the transport of early endosomes and hitchhiking cargoes such as peroxisomes (Abenza et al., 2009; Christensen and Reck-Peterson, 2022; Egan et al., 2012; Guimaraes et al., 2015; Lenz et al., 2006; Müntjes et al., 2021; Plamann et al., 1994; Qiu et al., 2019; Salogiannis et al., 2021; Salogiannis et al., 2016; Xiang et al., 1994; Xiang et al., 1995; Zekert and Fischer, 2009). As a defect in dynein or LIS1 function causes early endosomes to accumulate abnormally near the hyphal tip, early-endosome distribution is an excellent readout of dynein or LIS1 function (Lenz et al., 2006). Compared to dynein-mediated nuclear distribution, early-endosome distribution in *A. nidulans* appears to be even more sensitive to dynein abnormalities (Tan et al., 2014). However, in both the *ubaB*^Q247^* and Δ*sumO* mutants, we have detected no defect in the distribution of early endosomes labeled by mCherry-Rab5A (formerly called RabA) (Pinar and Peñalva, 2021). Specifically, none of the mutants showed any obvious hyphal-tip accumulation of early endosomes (Figure 7), in contrast to a recently studied kinesin-1 mutant defective in dynein-mediated early endosome transport (Qiu et al., 2023b). These results indicate that SUMOylation does not play an important role in dynein or LIS1 function in *A. nidulans*.

**Figure 7.**
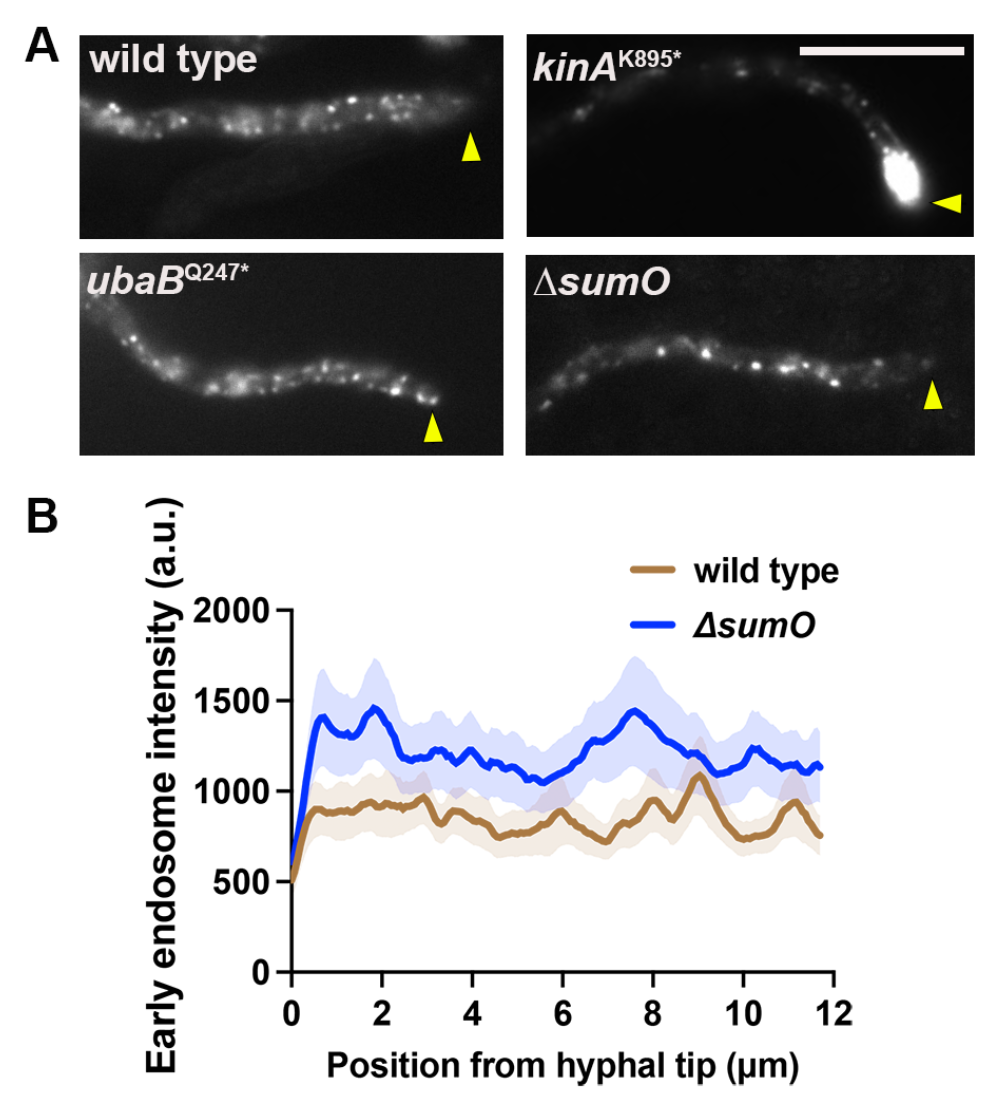
Loss of SUMOylation does not cause early endosomes to accumulate abnormally at the hyphal tip. (A) Distribution of mCherry-Rab5A-labeled early endosomes in the wild type, the *ubaB*^Q247^* mutant and the Δ*sumO* mutant. The hyphal tip is indicated by a yellow arrowhead. The *kinA*^K895*^ mutant was used as a control (Qiu et al., 2023b), as most hyphal tips of this mutant show a clear accumulation of mCherry-Rab5A-labeled early endosomes, which indicates a defect in dynein-mediated transport of early endosomes from the hyphal tip. The *ubaB*^Q247^* and Δ*sumO* mutants show a normal distribution of early endosomes, as compared to the wild type and the *kinA*^K895*^ mutant. Bar, 10 μm. (B) Line scans of mCherry-Rab5A (early endosomes) fluorescence intensity in the wild type and the Δ*sumO* mutant. XY graphs with mean (solid lines) and standard error (SEM, shading) were generated by Prism 9. The overall intensity of mCherry-Rab5A appeared higher in the mutant than in the wild type along the hyphae but the differences were not statistically significant (p>0.9999 for all the points from 0 to 11.700 μm from the hyphal tip with 0.065 μm as intervals, two-way ANOVA with Bonferroni’s multiple comparisons test, n=31 hyphae for both the wild-type control and the Δ*sumO* mutant).

Given the different requirements for SUMOylation in different cell types, whether SUMOylation plays a role in LIS1 regulation still needs to be tested in other cell types. This is especially true for *S. cerevisiae* where the interactions between the LIS1 homolog PAC1 (Geiser et al., 1997) and proteins in the SUMOylation pathway was first revealed (Alonso et al., 2012). Pac1/LIS1 binds dynein directly and affects dynein in various ways (Gillies et al., 2022; Markus et al., 2020). *A. nidulans* and *S. cerevisiae* LIS1 homologs help to overcome an autoinhibited dynein conformation since the phi mutations preventing the autoinhibited state (Zhang et al., 2017) partially compensate for LIS1 deficiency (Marzo et al., 2020; Qiu et al., 2019), while mammalian LIS1 enhances the in vitro assembly of activated dynein-dynactin-cargo adapter complex (Elshenawy et al., 2020; Htet et al., 2020). Intriguingly, prior to being offloaded with dynactin to the cortex to bind the cortical adapter Num1 (Omer et al., 2018), dynein is accumulated at the microtubule plus end in a Pac1/LIS1-dependent but dynactin-independent fashion (Lee et al., 2003; Sheeman et al., 2003). This differs from *A. nidulans* and *Ustilago maydis* where the plus-end dynein accumulation depends on dynactin but not LIS1 (Lenz et al., 2006; Zhang et al., 2003) and from mammalian cells (or in vitro with mammalian proteins) where both LIS1 and dynactin are important for the plus-end-targeting of dynein (Baumbach et al., 2017; Jha et al., 2017; Splinter et al., 2012). Given these differences, it is conceivable that there are differences in how LIS1 is regulated among various organisms and cell types.

## Materials and Methods1

### *A. nidulans* strains and media

*A. nidulans* strains used in this study are listed in Table 1. Detailed medium compositions can be found in a recent publication (Qiu et al., 2023a). Solid rich medium was made of either YAG (0.5% yeast extract and 2% glucose with 2% agar) or YAG+UU (YAG plus 0.12% uridine and 0.11% uracil). Solid minimal medium containing 1% glucose was used for selecting progeny from a cross. For live cells imaging experiments, cells were cultured in liquid minimal medium containing 1% glycerol for overnight at 32°C or containing 1% glucose for overnight at room temperature.

### Live cell imaging and analyses

Microscopic images used in Figures 1, 2, 5, 7, S2, S3 and S4 were generated using a Nikon Ti2-E inverted microscope with Ti2-LAPP motorized TIRF module and a CFI apochromat TIRF 100 x 1.49 N.A. objective lens (oil). The microscope was controlled by NIS-Elements software using 488 nm and 561 nm lines of LUN-F laser engine and ORCA-Fusion BT cameras (Hamamatsu). For all images, cells were grown in the LabTek Chambered #1.0 borosilicate coverglass system (Nalge Nunc International, Rochester, NY). Images were taken at room temperature. All the images were taken with a 0.1-s exposure time (binning: 2×2). For line scans presented in Figure 7, we draw a line starting from the hyphal tip in the middle of the hypha and get the average intensity value for the width of 2 mm, which is normally the hyphal width. For Figures 3 and 4, and Video 1, live-cell-imaging was performed as previously described (Bieger et al., 2021). Specifically, it was performed on a Nikon Ti-E Eclipse inverted epifluorescence microscope, equipped with a Perfect Focus System (Nikon) and a motorized Piezo stage, using a 100x 1.49 N.A. oil immersion Apo TIRF Nikon objective. GFP and DsRed were excited using an AURA II triggered illuminator with 485nm and 560nm LEDs, respectively, and detected using a Zyla 4.2 sCMOS camera (Andor, Oxford Instruments). All hardware was controlled by NIS-Elements Advanced Research (version 4.60). Two-color 3D time lapse data sets were deconvolved, with spherical aberration correction and background subtraction, using the “Automatic” 3D deconvolution option in NIS-Elements Advanced. Video 1 was generated in Imaris (9.5.1; Bitplane). Images used in Figure 6 were captured using an Olympus IX73 inverted fluorescence microscope linked to ORCA-FLASH4.0LT+ SCMOS CAMERA and controlled by cellSens Dimension Version 3 software (Olympus Inc.). An UPlanSApo 100× objective lens (oil) with a 1.40 numerical aperture (NA) was used. An ET-EGFP/mCherry Dual Multiband Filter Set w/BX3 cube (purchased from Olympus Inc.) was used.

### Mutagenesis and identification of the mutation that causes the chromatin bridge phenotype

UV mutagenesis was done as previously described (Qiu et al., 2023b). For identifying the causal mutation, we first crossed the identified mutant to a wild-type strain. We then picked five mutant progenies from the cross with the same colony phenotype, and picked five wild-type progeny (Note that the chromatin-bridge phenotype of the mutant is linked to the colony phenotype of poor conidiation). The spores of the five progeny in each group were mixed. Both the mutant and wild-type spores were cultured to make genomic DNA using the QIAGEN DNeasy Plant Mini Kit. The mutant and wild-type genomic DNAs were then sent to Otogenetics Corp for whole genome sequencing. By comparing the whole genome sequencing data from the two groups, we identified 1248 sequence variants in the mutant group but not in the wild type. We further annotated these variants in MySQL database using *Aspergillus nidulans* gene features table downloaded from (http://www.aspergillusgenome.org/download/chromosomal_feature_files/A_nidulans_FGSC_A 4/), a site that is no longer available and has been replaced by FungiDB (https://fungidb.org/fungidb/app). We identified 140 sequence variants positioned in the coding regions of 48 genes. We then manually checked these 48 genes using the IGV software and identified the single mutation in AN2450, which encodes UbaB.

### Construction of the **Δ***ubaB* mutant

For constructing the Δ*ubaB* mutant, we first made the ΔUbaB construct with the selective marker *pyrG* from *Aspergillus fumigatus*, *AfpyrG*, in the middle of the linear construct (Figure S1). Specifically, we used APYRGF (5’-TGCTCTTCACCCTCTTCGCG-3’), and APYRGR (5’-CTGTCTGAGAGGAGGCACTGA-3’) as primers and the pFNO3 plasmid (Yang et al., 2004) as template to amplify a 1.9-kb AfpyrG fragment. We used UBABNN1 (5’- CCGCTTGCTATAAGATTTACGATC-3’) and UBABNC (5’-CGCGAAGAGGGAGAAGAGCAAGTGCGATCAATAGTCAATAAAGC-3’) as primers and wild-type genomic DNA as template to amplify a 1.1-kb fragment upstream of the UbaB coding sequence. We used UBABCN (5’- ATCAGTGCCTCCTCTCAGACAGTAACGCACGCATATGCGCA-3’) and UBABCC1 (5’-CAACCAGTCGACATTCTCG-3’) as primers and wild-type genomic DNA as template to amplify a 1.1-kb fragment downstream of the UbaB coding sequence. We then used 2 primers: UBABNN (5’- CGCAAGCTCTATTTGTGCGAC-3’) and UBABCC (5’-GCAGCAGGCATTTCAACATCC −3’) to perform a fusion PCR to fuse the three fragments and obtained a 3.9-kb fragment, which we transformed into the *A. nidulans* strain XX357 containing Δ*nkuA*. Several transformants were obtained that show a “small-colony” phenotype. Homologous integration of the deletion construct was confirmed by PCR using the following two pairs of primers: UBABNN1 and AFpyrG3 (5’-GTTGCCAGGTGAGGGTATTT-3’); UBABCC1 and AfpyrG5 (5’-AGCAAAGTGGACTGATAGC-3’).

### Construction of a strain containing the *ubaB-GFP* allele at the *ubaB* locus

The construction was done by using standard procedures used in *A. nidulans* (Nayak et al., 2006; Szewczyk et al., 2006; Yang et al., 2004). For constructing the UbaB-GFP fusion, we used the following six oligos to amplify genomic DNA and the GFP-*AfpyrG* fusion (Yang et al., 2004): UBABGNC (5’-GGCTCCAGCGCCTGCACCAGCTCCGTCATCATCGATCAGAATAGCGC-3’), UBABGNN (5’-CACTGCAGTCCGAAGAAAGC-3’), UBABCN (5’-ATCAGTGCCTCCTCTCAGACAGTAACGCACGCATATGCGCA-3’), UBABCC1, GAGAF (5’- GGAGCTGGTGCAGGCGCTG-3’), and pyrG3 (5’-CTGTCTGAGAGGAGGCACTGAT-3’). Specifically, UBABGNC and UBABGNN were used to amplify the 1-kb fragment in the codon region, and UBABCN and UBABCC1 were used to amplify the 1.1-kb fragment in the 3’ untranslated region, using wild-type genomic DNA as template, and GAGAF and PyrG3 were used to amplify the 2.7-kb GFP-AfpyrG fragment using the pFNO3 plasmid DNA as template (Yang et al., 2004). We then used two oligos, UBABGNN1 (5’-GGTTCTGGTGTTCGACAAGG- 3’) and UBABCC for a fusion PCR of the three fragments to generate the UbaB-GFP-AfpyrG fragment that we used to transform into a wild-type strain containing the Δ*nkuA* allele (Nayak et al., 2006). The transformants were then screened by microscopically observing the GFP signals in the cells, and the homologous integration of the fusion DNA into the endogenous UbaB locus was confirmed by PCR. Specially, a 1.1-kb product was obtained with the two following oligos: AfpyrG5 and UBABCC1.

## Data availability Statement

Strains used and created in this work (listed in Table S1) are available upon request.

## Acknowledgements

We thank Stephen Osmani and Miguel Peñalva for Aspergillus strains, the Fungal Genetic Stock Center for the pFNO3 plasmid, and Stephen Osmani for depositing it. We thank David Pellman for very helpful discussions and Michael Lichten for help with early literature on yeast sumo mutants. This work was funded by the National Institutes of Health R35GM140792 (to X.X.) and NIH grant 1R15GM132869-01 (to M.J.E.) and the Irving S. Johnson Fund of the University of Kansas Endowment (to B.R.O.). Disclaimer: The opinions and assertions expressed herein are those of the author(s) and do not necessarily reflect the official policy or position of the Uniformed Services University or the Department of Defense.

## Online Supplemental Materials

**Table S1.** *Aspergillus nidulans* strains used in this study

| **Strain** | **Genotype** | **Source** |
| --- | --- | --- |
| TNO2A3 | $\Delta nkuA::argB$; *pyrG89*; *pyroA4* | (Nayak et al., 2006) |
| JZ850 | *ubaB*-GFP-*Afp**pyrG*; $\Delta nkuA::argB$; *pyrG89* | This work |
| JZ856 | $\Delta ubaB$-*Afp**pyrG*; *hhoA*-GFP-*AfriboB*; *argB2::[argB\*-alcAp::mCherry-RabA]*; $\Delta nkuA::argB$; *pyrG89*; *pantoB100* | This work |
| XX357 | *hhoA*-GFP-*AfriboB*; *argB2::[argB\*-alcAp::mCherry-RabA]*; $\Delta nkuA::argB$; *pyrG89*; *pantoB100* | This work |
| XX402 | *hhoA*-GFP-*AfriboB*; *argB2::[argB\*-alcAp::mCherry-RabA]*; $\Delta nkuA::argB$; *pantoB100* | This work |
| XX429 | *nimT23*; *hhoA*-GFP-*AfriboB*; *pyrG89*; possibly $\Delta nkuA::argB$ | This work |
| XX521 | *ubaB*<sup>Q247*</sup>; *hhoA*-GFP-*AfriboB*; *yA::NLS-DsRed*; possibly $\Delta nkuA::argB$ | This work |
| XX522 | *ubaB*<sup>Q247*</sup>; *hhoA*-GFP-*AfriboB*; *argB2::[argB\*-alcAp::mCherry-RabA]*; possibly $\Delta nkuA::argB$ | This work |
| XX523 | *hhoA*-GFP-*AfriboB*; *argB2::[argB\*-alcAp::mCherry-RabA]*; possibly $\Delta nkuA::argB$ | This work |
| XX541 | $\Delta ubaB$-*Afp**pyrG*; *nimT23*; *hhoA*-GFP-*AfriboB*; *pyrG89*; possibly $\Delta nkuA::argB$ | This work |
| XX554 | $\Delta sumO$-*Afp**pyrG*; *hhoA*-GFP-*AfriboB*; possibly $\Delta nkuA::argB$; possibly *pyrG89* | This work |
| XX790 | *hhoA*-GFP-*AfriboB*; $\Delta yA::NLS-DsRed$; possibly $\Delta nkuA::argB$ | This work |
| XX799 | *ubaB*-GFP-*Afp**pyrG*; $\Delta yA::NLS-DsRed$; possibly $\Delta nkuA::argB$; possibly *pyrG89* | This work |
| XX804 | $\Delta ubaB$-*Afp**pyrG*; *hhoA*-GFP-*AfriboB*; *argB2::[argB\*-alcAp::mCherry-RabA]*; *pabaA1*; *yA2* | This work |
| XX900 | $\Delta ubaB$-*Afp**pyrG*; *hhoA*-GFP-*AfriboB*; *yA::NLS-DsRed*; possibly $\Delta nkuA::argB$; possibly *pyrG89* | This work |
| XX902 | $\Delta ubaB$-*Afp**pyrG*; GFP-*tubA*; *yA::NLS-DsRed*; possibly $\Delta nkuA::argB$; possibly *pyrG89* | This work |
| XX903 | $\Delta sumO$-*Afp**pyrG*; *hhoA*-GFP-*AfriboB*; *argB2::[argB\*-alcAp::mCherry-RabA]*; possibly $\Delta nkuA::argB$; possibly *pyrG89* | This work |
| XX906 | *nimT23*; *hhoA*-GFP-*AfriboB*; possibly $\Delta nkuA::argB$ | This work |

**Figure S1.**
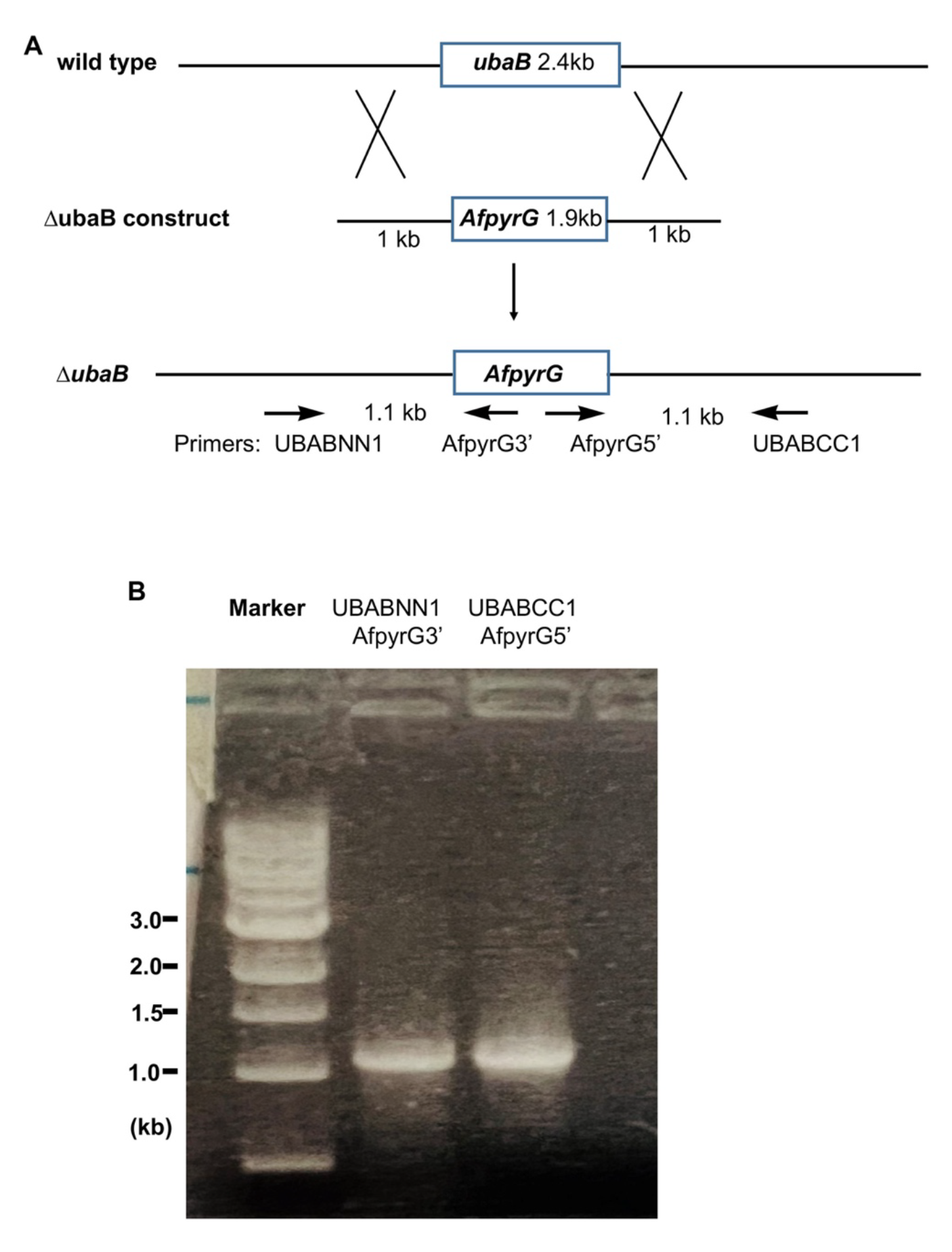
Site-specific integration of the ΔubaB construct into the genome. (A) A diagram showing the wild-type allele of the *ubaB* locus (wild-type), the ΔubaB linear construct with the *AfpyrG* marker flanked by the 5’ and 3’ flanking sequences of the *ubaB* gene, and the Δ*ubaB* allele (Δ*ubaB*). Homologous recombination events between the ΔubaB construct and the wild-type genome (wild type) are indicated by crosses. The resulting Δ*ubaB* allele is shown at the bottom. The positions of the primers used for PCR analyses are indicated by arrows. Note that the primers UBABNN1 and UBABCC1 are located outside of the flanking sequences of the ΔubaB construct and thus the PCR reactions will produce the expected products of 1.1-kb only when the construct integrated into the *ubaB* locus. (B) Result of a PCR analysis on genomic DNAs from the Δ*ubaB* mutant. Image of a DNA agarose gel is shown, and primers are indicated at the top.

**Figure S2.**
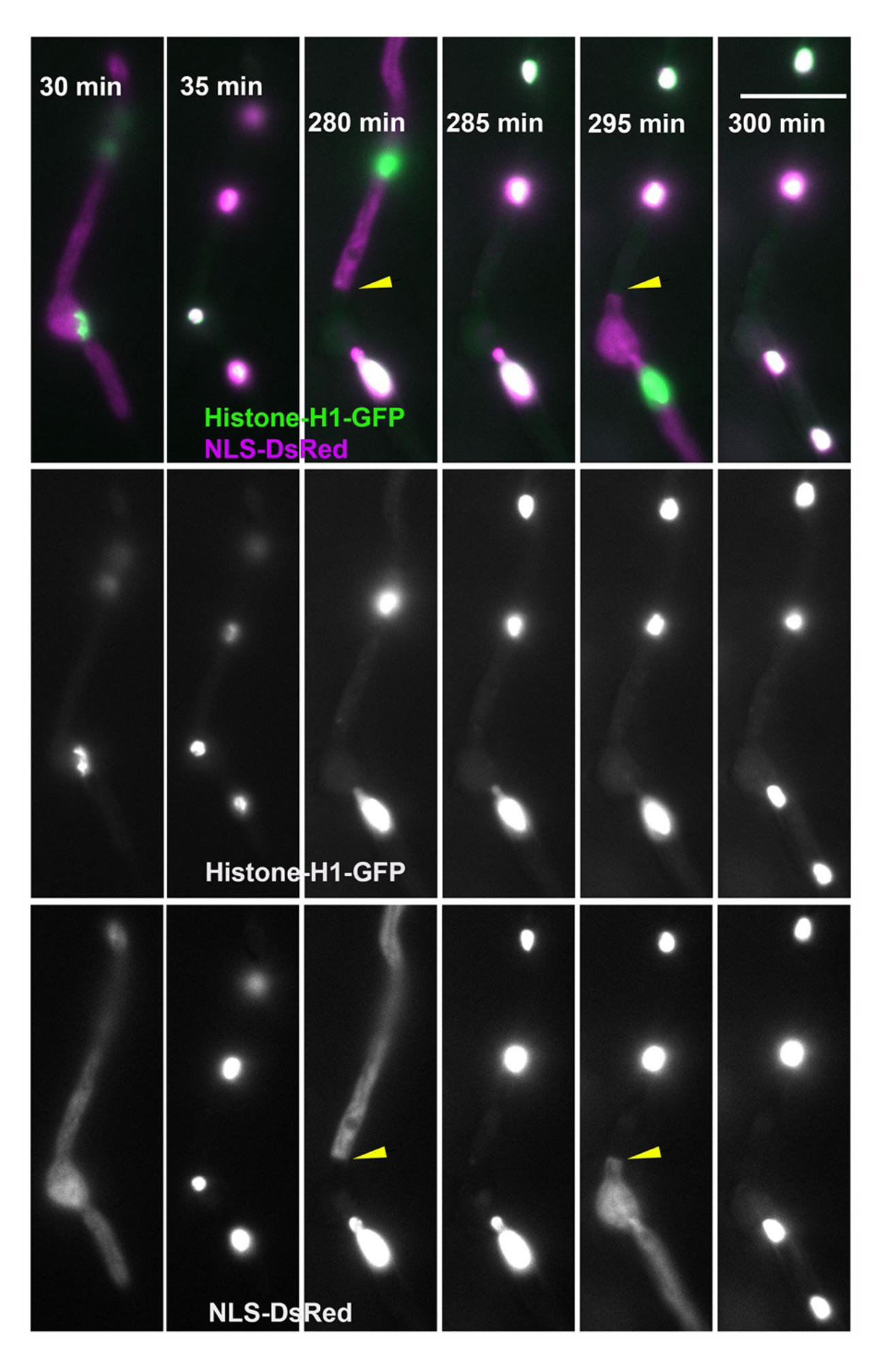
A wild-type hypha containing both Histone-H1-GFP and NLS-DsRed. Six different time points of a time-lapse sequence (taken at 5-min intervals) are presented to show that the NLS-DsRed fusion proteins become cytoplasmic during mitosis and that the position of a closed septum (yellow arrowhead) can be inferred from the boundary of the cytoplasmic NLS-DsRed signals (Shen et al., 2014). At 30 min, the nuclei were undergoing mitosis as indicated by the cytoplasmic NLS-DsRed signals. At 35 min, daughter nuclei at G1 (labeled by NLS-DsRed signals) were formed. The 280-min time point shows the position of a closed septum (yellow arrowhead), as the upper compartment contains cytoplasmic NLS-DsRed signals and the lower compartment contains an interphase nucleus at G2. At 285 min, two daughter nuclei formed after mitosis in the upper compartment. At 295 min, the G2 nucleus in the lower compartment entered mitosis. At 300 min, two daughter nuclei were formed after mitosis in the lower compartment. Bar, 10 μm.

**Figure S3.**
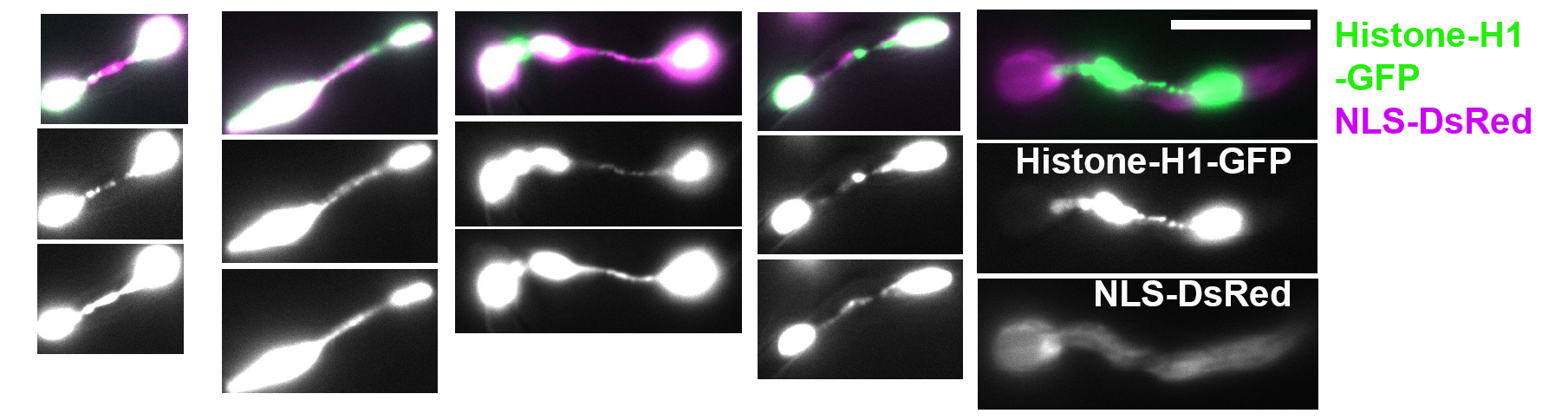
Multiple images of the Δ*ubaB* mutant containing both Histone-H1-GFP (middle) and NLS-DsRed (lower) (note that the merge is shown on top). Most chromatin bridges can enter interphase as evidenced by the nuclear NLS-DsRed signals. Note that on the right side, the two nuclei connected by a chromatin bridge are inside a mitotic cell, as evidenced by cytoplasmic NLS-DsRed signals. Bar, 10 μm.

**Figure S4.**
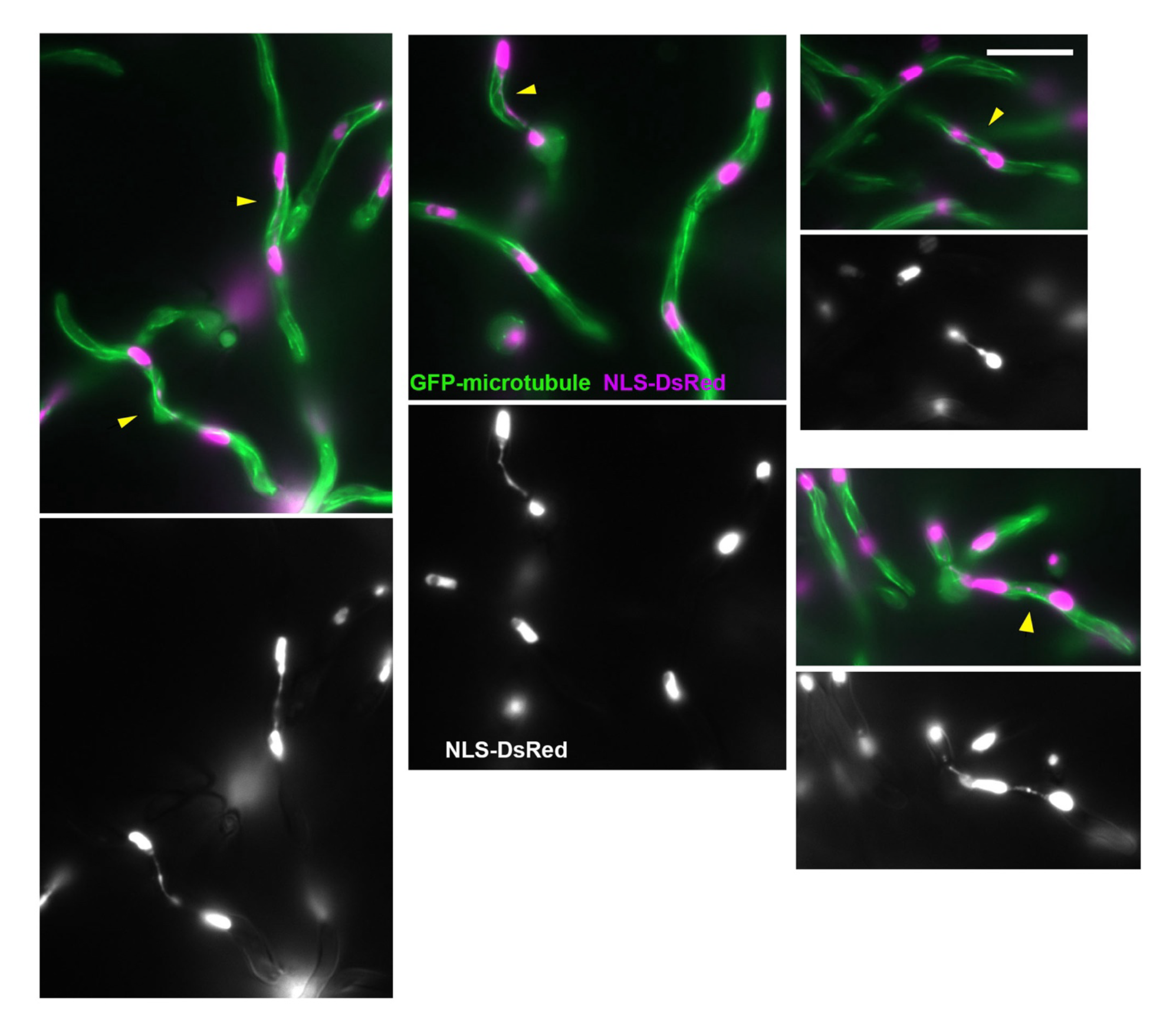
Multiple mages of the Δ*ubaB* mutant containing GFP-microtubules and NLS-DsRed (lower) (note that the merge is shown on top). Most NLS-DsRed-labeled bridges (indicated by yellow arrows on the merged images) can be seen to be surrounded by cytoplasmic microtubules typical of interphase cells. A bridge in the lower right panel (indicated by a bigger yellow arrow) may possibly be connected by an elongated spindle (although this could still be debated because a typical mitotic spindle has better defined poles at the two ends) (Figure 2B) (Chemudupati et al., 2019; Horio and Oakley, 2005; Li et al., 2005). Bar, 10 μm.

**Video 1.** An 11-hour time-lapse sequence with 10-min intervals showing a *ubaB*^Q247^* mutant hypha whose nuclei are labeled with Histone-H1-GFP and NLS-DsRed. At the time point 1:39:58.317, the presence of a closed septum became obvious as evidenced by the appearance of NLS-DsRed signals only on the lower side of the chromatin bridge. This video also shows that the hypha entered mitosis at 6:39:58.991 and exited from mitosis at 6:59:59.462 as indicated by the disappearance and reappearance of NLS-DsRed signals in the nuclei, while the chromatin bridges were not resolved. Micrographs represent maximum intensity projections of two-color Z-series acquired 0.2 μm intervals spanning the entire depth of the hyphae.

